# Iron chelation by oral deferoxamine treatment decreased brain iron and iron signaling proteins

**DOI:** 10.1101/2024.08.14.607970

**Authors:** Max A. Thorwald, Jose A. Godoy-Lugo, Naomi S. Sta Maria, Ararat Chakhoyan, Peggy A. O’Day, Russell E. Jacobs, Berislav Zlokovic, Caleb E. Finch

**Affiliations:** Leonard Davis School of Gerontology, University of Southern California, Los Angeles, CA; Zilkha Neurogenetic Institute, Keck School of Medicine, University of Southern California, Los Angeles CA; Department of Physiology and Neuroscience, Keck School of Medicine, University of Southern California, Los Angeles CA; Life and Environmental Sciences Department, University of California, Merced, CA; Dornsife College, University of Southern California, Los Angeles, CA

**Keywords:** Alzheimer’s disease, iron, DFO, MRI, APP, secretase enzymes, lipid peroxidation

## Abstract

**Background:** Deferoxamine (DFO) and other iron chelators are clinically used for cancer and stroke. They may also be useful for Alzheimer’s disease (AD) to diminish iron from microbleeds. DFO may also stimulate antioxidant membrane repair which is impaired during AD. DFO, and other chelators do enter the brain despite some contrary reports.

**Objective:** Low dose, oral DFO was given in lab chow to wildtype (WT) C57BL/6 mice to evaluate potential impact on iron levels, iron-signaling and storage proteins, and amyloid precursor protein (APP) and processing enzymes. Young WT mice do not have microbleeds or disrupted blood-brain barrier of AD mice.

**Methods:** Iron was measured by MRI and chemically after two weeks of dietary DFO. Cerebral cortex was examined for changes in iron metabolism, antioxidant signaling, and APP processing by Western blot.

**Results:** DFO decreased brain iron by 18% (MRI) and decreased seven major proteins that mediate iron metabolism by at least 25%. The iron storage proteins ferritin light and heavy chain decreased by at least 30%. APP and secretase enzymes also decreased by 30%.

**Conclusions:** WT mice respond to DFO with decreased APP, amyloid processing enzymes, and antioxidant repair. Potential DFO treatment for early-stage AD by DFO should consider the benefits of lowered APP and secretase enzymes.

## INTRODUCTION

Iron is the most abundant transition state metal and critical body-wide for oxygen binding by heme, electron transfer by mitochondria, and as a cofactor for DNA synthesis and repair. Other transition state metals including zinc, copper, aluminum, and others are also essential. Toxic excess of these metals is implicated in Alzheimer’s disease (AD), through increased lipid peroxidation in humans (1,2) and AD mouse models (3,4). Of brain total fats, more than half are poly-unsaturated fatty acids that are readily oxidized by these transition state metals.

Cerebral microbleeds (MBs) during AD contribute to increased brain iron (4,5). ‘Naked’ MBs, those without amyloid plaque localization, arise by age 2 months in the ApoEFAD mouse model (5): at 4-6 months MBs and extracellular fibrillar β-amyloid (Aβ) plaques have become colocalized, suggesting an interaction between these pathologies. The nascent MBs are independent of cerebral amyloid angiopathy in EFAD mice, which arise in different cortical layers (5). Fibrillar amyloid resides in areas of high iron content by MRI in individuals with mild cognitive impairment (6). We (4) and others (7,8) have analyzed relationships of brain iron to regional specificity in AD. The cerebellum which has minimal AD neurodegeneration has lower iron mediated lipid peroxidation and neuron atrophy.

Iron chelators are potential AD therapeutics. The chelator deferoxamine (DFO) was originally discovered in 1960 and soon found to bind transition state metals aluminum, copper, zinc with high affinity (10–12). DFO was the first FDA approved iron chelator in 1968 and is still prescribed for thalassemia and dialysis (13,14). DFO has been used in 103 clinical trials, mostly for arthritis, cancer, and diabetes. Of these, only 7 trials were related to hemorrhage and none for the treatment of neurological diseases. The first and only clinical trial of DFO for AD in 1991 showed slowed cognitive decline over a 24-month period (15). Although DFO’s half-life is just 10 minutes in plasma (16), two studies with DFO showed its retention by brain tissues after intravenous (i.v.) injection in dogs by radio-labelling (17) and by intra-peritoneal (i.p.) injection in rats (18). These decades old reports are not cited by some which state that DFO does not cross the blood brain barrier (BBB) (19) or crosses “poorly”, which would limit efficacy (20,21). Toxicity of iron chelators at high doses is another concern (12), e.g. ocular neuropathy for 50 mg/kg/day.

Mouse AD models have shown DFO benefits in several labs. APP/PS1 mice with different DFO administration and duration: intranasally (i.n.) for 18 weeks (22) or i.p. 1 week (23). Both treatments showed potential benefits to AD, with a 30% decrease in soluble Aβ peptides. Moreover, iron was decreased by 50% (Prussian blue histochemistry), and short term spatial memory (water maze) was improved (23). Recently, we showed major impact of DFO on ApoEFAD mice with decreased fibrillar Aβ and lipid peroxidation; DFO in diet and i.p. caused similar responses. DFO increased levels of the transcription factor Nrf2 and several antioxidants it transcriptionally regulates (4). For wildtype mice, others showed improved memory for DFO administered by i.n., but not i.p (22). DFO also increased Nrf2 and its targets in other cell types, e.g. murine chondrocytes (24) and in cancer cells but with increased oxidative damage (25). In an ALS model DFO by i.p. has also been shown to lower iron in the lumbar anterior horn slowing progression of motor-neuron injury (26). The present study expands these findings to responses of wildtype mice to DFO for brain iron, iron signaling, antioxidant pathways, and amyloid precursor protein (APP) and its processing enzymes. MRI is introduced for measurement of mouse brain iron.

## RESULTS

### Deferoxamine (DFO) decreases total brain iron, but not in AD relevant regions of frontal cortex or hippocampus

Young wildtype (WT) mice were fed DFO in lab chow 10mg/kg/day for two weeks, starting at 5.5 months. Brain region responses to DFO for total iron were analyzed by three techniques: magnetic resonance imaging (MRI) (**Fig. 1A**) and chemically by **D)** inductively coupled mass spectrometry (ICP-MS) and **E)** for heme.

**Figure 1:**
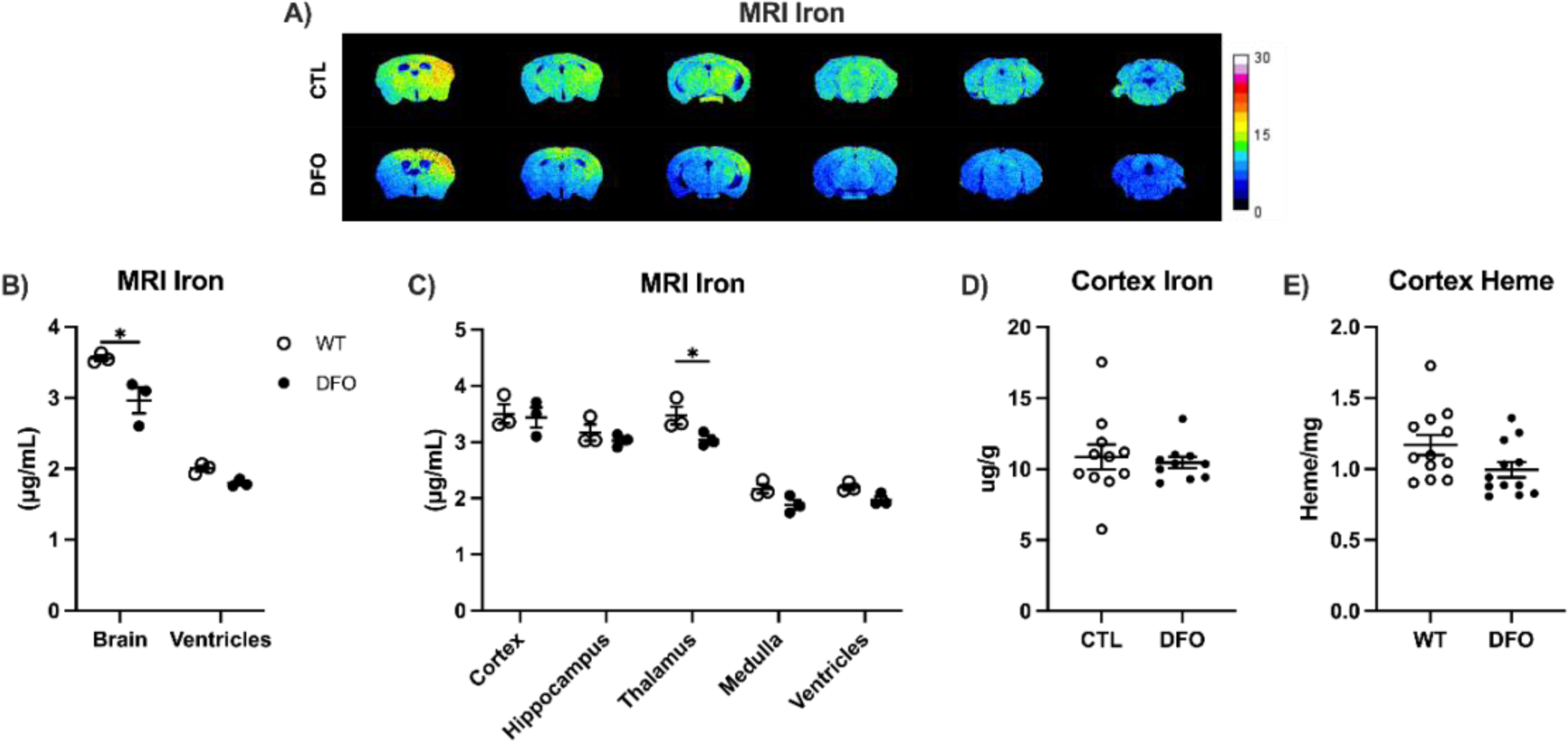
DFO decreased total brain iron with brain region specificity. **A)** Representative sequential R_2_ mapping of coronal slices from multi-echo multi-slice (MEMS) MRI for total iron. Iron levels were calibrated by standard curve with phantom containing iron oxide particles (**S.**Fig. 1A). Extrapolated assayed by **D)** inductively coupled mass spectrometry (ICP-MS) and **E)** tissue heme. Statistics: two-way ANOVA with Tukey’s posthoc for between group comparisons (**B,C;** WT vs DFO only) by brain region; two-tailed t-test **(D,E)**: *p<0.05; for full statistical comparisons, **S.**Fig. 1.

By MRI, DFO lowered whole brain iron 18% (**Fig. 1A**). DFO did not alter iron in cerebral cortex, hippocampus, or medulla (**Fig. 1C**), whereas thalamus was 15% lower (**Fig. 1C**). Iron in lateral ventricles was unaltered, implying that DFO did not alter intraventricular fluid iron (**Fig. 1B**). Brain region iron also differed between coronal sections of olfactory bulb, anterior (-Bregma), middle (+Bregma), and cerebellum. The anterior portion showed 19% less iron (**S.Fig. 1B,C**). Total iron by ICP-MS was unchanged by DFO in cerebral cortex, confirming the MRI findings (**Fig. 1D**). Because DFO chelates aluminum, copper, and zinc (11), these were assayed but did not respond to DFO (**S.Fig. 2**). Heme was also not altered by DFO in cerebral cortex (**Fig. 1E**).

**Figure 2:**
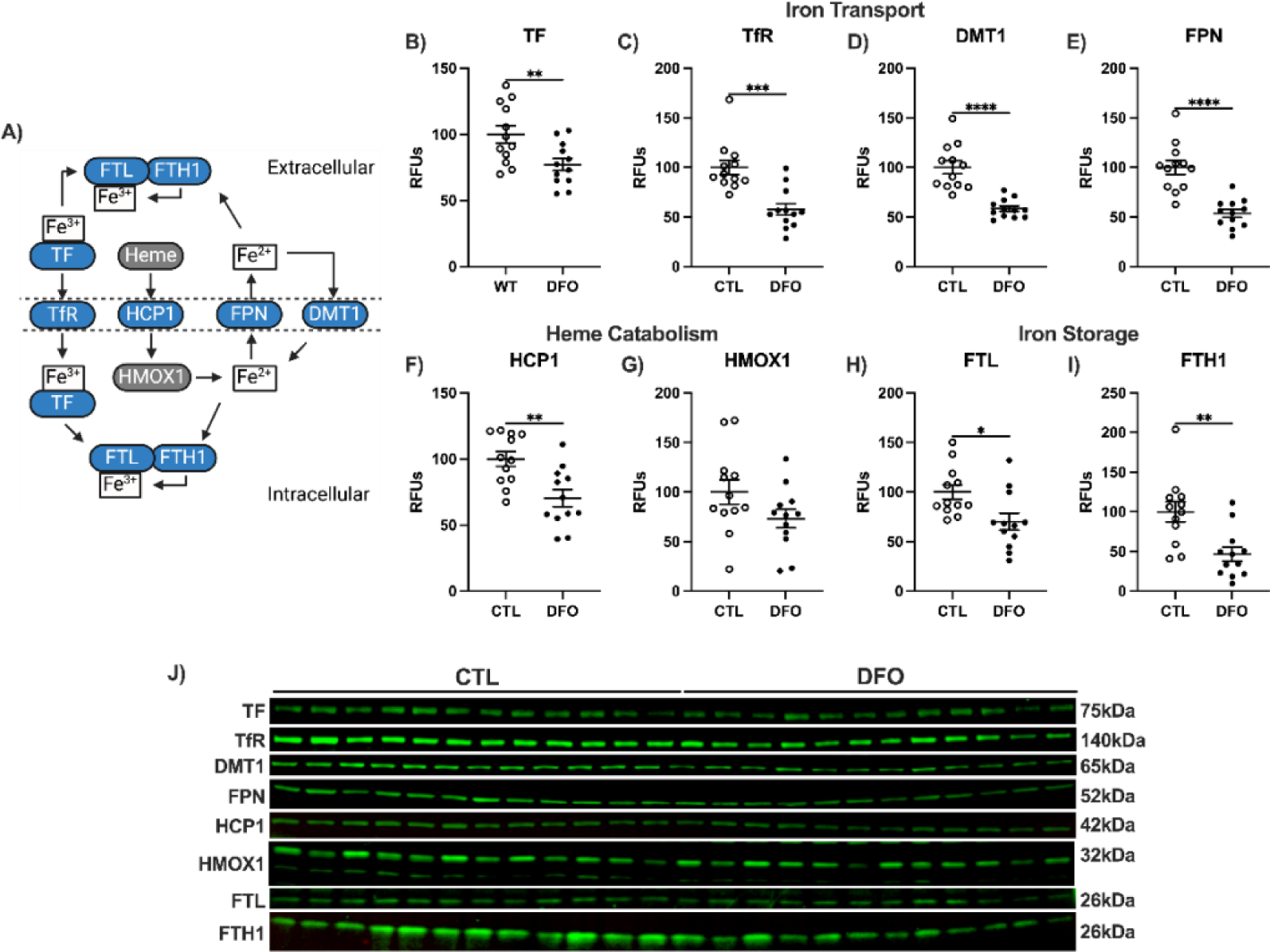
Iron-signaling proteins in cerebral cortex are decreased by DFO. **A)** Schematic for iron transport protein to DFO: blue, increase; grey, no change. **B-I** Western blot, relative fluorescent units (RFUs) for TF, TfR, DMT1, FPN, HCP1, HMOX1, FTL, FTH1. **J**) representative Western blots. Two-tailed t-test: *p<0.05, **p<0.01, ***p<0.001, ****p<0.0001.

### DFO decreased iron signaling and transport proteins in cerebral cortex

Circulatory iron is transported by transferrin (TF) as ferric iron (Fe^3+^) and imported in part by the transferrin receptor (TfR). Ferrous iron (Fe^2+^) also enters brain cells by divalent metal transporter 1 (DMT1). Ferrous iron is generated intracellularly from heme breakdown by heme-oxygenase 1 (HMOX1) after import by heme carrier protein 1 (HCP1). Ferrous iron is then oxidized to ferric iron by ferroxidase within ferritin heavy chain 1 (FTH1). Normal and pathological iron is stored on complexes of ferritin light chain (FTL) in heterogenous cage structures with up to 4,500 Fe^3+^ atoms (27). Export of iron is mediated solely by ferroportin (FPN) as Fe^2+^ or by lysosomal degradation of ferritin complexes (28) (**Fig. 2A**).

Cerebral cortex was examined because of its neurodegeneration in early stages of AD. Despite the lack of DFO impact on total iron (**Fig.1C,**), DFO decreased many iron import proteins: TF, TfR, DMT1 decreased by 25% or greater (**Fig. 2B-D**); iron export protein FPN by -45% (**Fig. 2E**); heme catabolism protein HCP1 decreased by 30% (**Fig. 2F**). Iron storage proteins FTL and FTH1 were decreased by 30% (**Fig. 2H,I**). Only HMOX1 was unaltered by DFO (**Fig. 2G**). The decrease in iron signaling proteins without reductions in total cortex iron suggests an underlying complexity that DFO alters subcellular iron with brain region differences.

### Amyloid precursor protein (APP) and its processing enzymes are decreased by DFO

The amyloid precursor protein APP and the secretase enzymes were examined for selectivity of response to DFO. APP has multiple isoforms in most brain cells, including APP_695_, APP_751_, and APP_770_ that are generated by alternative splicing. APP is linked to iron through the regulation of its transcript by iron responsive proteins (29). DFO decreased APP protein by 25% (**Fig. 3A**). Neuronal APP_695_ is cleaved by two secretase enzymes: the α-secretase ADAM10 (A Disintegrin and metalloproteinase domain-containing protein 10), or by β-secretase 1 (BACE1).

**Figure 3:**
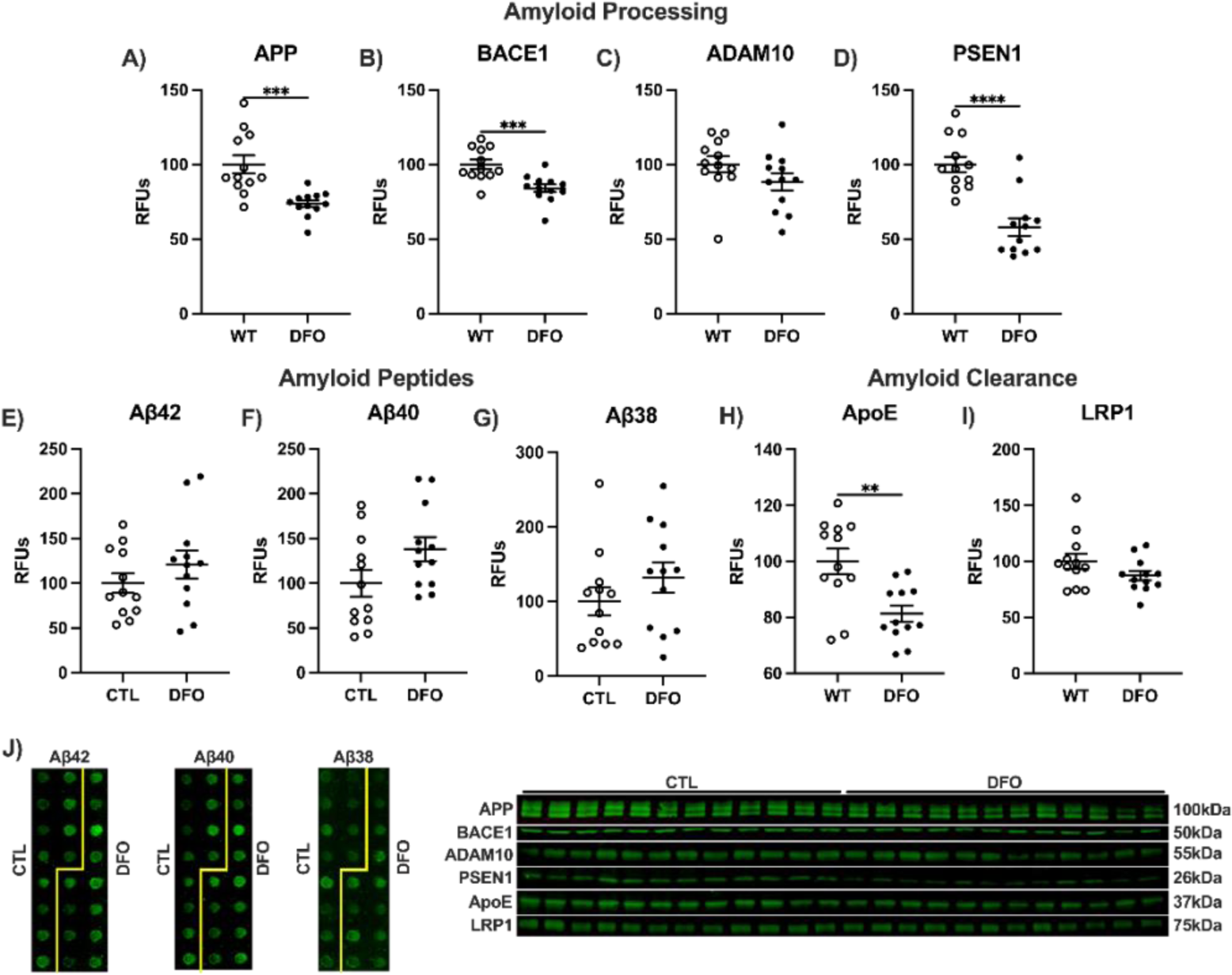
DFO selectively decreased APP and secretase levels, but did not alter the main Aβ peptides, shown as dot or Western blots as relative fluorescent units (RFUs) for **A)** APP, **B)** BACE1, **C)** ADAM10, **D)** PSEN1, **E)** Aβ42, **F)** Aβ40, **G)** Aβ48, **H)** ApoE, and **I)** LRP1. **J)** Representative images for dot and Western blots. Two-tailed t-test: **p<0.01, ***p<0.001, ****p<0.0001.

DFO decreased BACE1 protein -15%, while ADAM10 remained unchanged with DFO (**Fig. 3B,C**). The non-amyloidogenic ADAM10 processing of APP yields peptide products soluble APPα (sAPPα) and c-terminal fragment 83 (CTF83). Amyloidogenic processing of APP occurs through BACE1 in endosomes containing lipid rafts yielding CTF99 and sAPPβ. CTF99 and CTF83 are both cleaved by the γ-secretase complex that includes presenilin 1 (PSEN1) in its catalytic domain which DFO decreased 40% (**Fig. 3D**). Cleavage of CTF99 by the γ-secretase yields the APP intracellular domain and monomeric Aβ38, -40, -42, while CTF83 can only form p3. None of the Aβ peptides were decreased by DFO, despite the lowering of the APP and secretase enzymes **Fig. 3E-G**). Soluble Aβ peptides are cleared by binding to apolipoprotein E (ApoE) increasing recognition to the low-density lipoprotein receptor-related protein 1 (LRP1) for transport across the BBB. DFO decreased ApoE protein by -20%, while LRP1 remained unchanged (**Fig. 3H-I**).

### Antioxidant pathways that mitigate lipid peroxidation respond minimally to DFO

Next, we examined antioxidant pathways for DFO response. Iron, copper, and other transition state metals initiate lipid peroxidation through Fenton chemistry. In our recent study of EFAD mice, DFO increased Nrf2, a transcription factor which regulates antioxidant genes and several target proteins (3). We extend these studies to WT mice. Also relevant to Nrf2 is the BTB domain and CNC homolog 1 (BACH1) which represses Nrf2 binding to these targets by competitive inhibition (30,31). Neither Nrf2 or BACH1 were altered by DFO in cerebral cortex, despite increased Nrf2 in EFAD mice (4) and in other non-brain cell (20,21) (**S.Fig. 3A,B**).

Glutathione-related proteins regulated by Nrf2 that generate GSH responded differentially to DFO. The critical antioxidant GSH is produced body-wide by glutamate cysteine ligase (GCL) and GSH synthetase. Cysteine and glutamate are ligated by the GCL heterodimer comprised of modifier (GCLM) and catalytic (GCLC) subunits. GCLM increases the rate of catalysis 4-fold for γ-glutamylcysteine formation and is the rate-limiting step in GSH biosynthesis (32). DFO decreased both GCLC and GCLM by 20% (**S.Fig. 3C,D**). Cysteine, rate-limiting for GSH synthesis, is imported as cystine by the Cystine/Glutamate transporter (xCT), which DFO did not alter (**S.Fig. 3E**).

Glutathione peroxidase 4 (GPx4), glutathione peroxidase 1 (GPx1), and peroxiredoxin 6 (Prdx6) are phospholipid hydroperoxidases that require GSH as a reductant. GPx4 and Prdx6 chemically reduce oxidized lipids and sterols within cell membranes (33,34). GPx4 was decreased by DFO, while GPx1 and Prdx6 were not altered (**S.Fig. 3F-H**). Ferroptosis suppressor protein 1 (FSP1) is GSH independent and reduces lipid radicals through the quinol cycle. FSP1 was also unaltered with DFO (**S.Fig. 3I**).

Lipid hydroperoxides that are not detoxified may decompose to reactive aldehydes such as 4-hydroxy-2-nonenal (HNE). Most free HNE is cleared through glutathione s-transferase family members, primarily GSTA4 in brain, or GSH independent aldehyde dehydrogenase 2 (ALDH2). GSTA4 was unchanged by DFO, while ALDH2 decreased 35% (**S.Fig. 3J,K**). Free HNE that is not cleared forms HNE adducts on proteins altering structure and function (35). DFO treatment did not alter HNE levels (**S.Fig. 3I**). Decreased xCT, GPx4, or increases in lipid peroxidation are all associated with iron mediated cell death ‘ferroptosis’ (36,37). **S.Fig. 3M** outlines antioxidants relevant to attenuating ferroptosis.

## Supplemental Figures

**Supplemental Figure 1:**
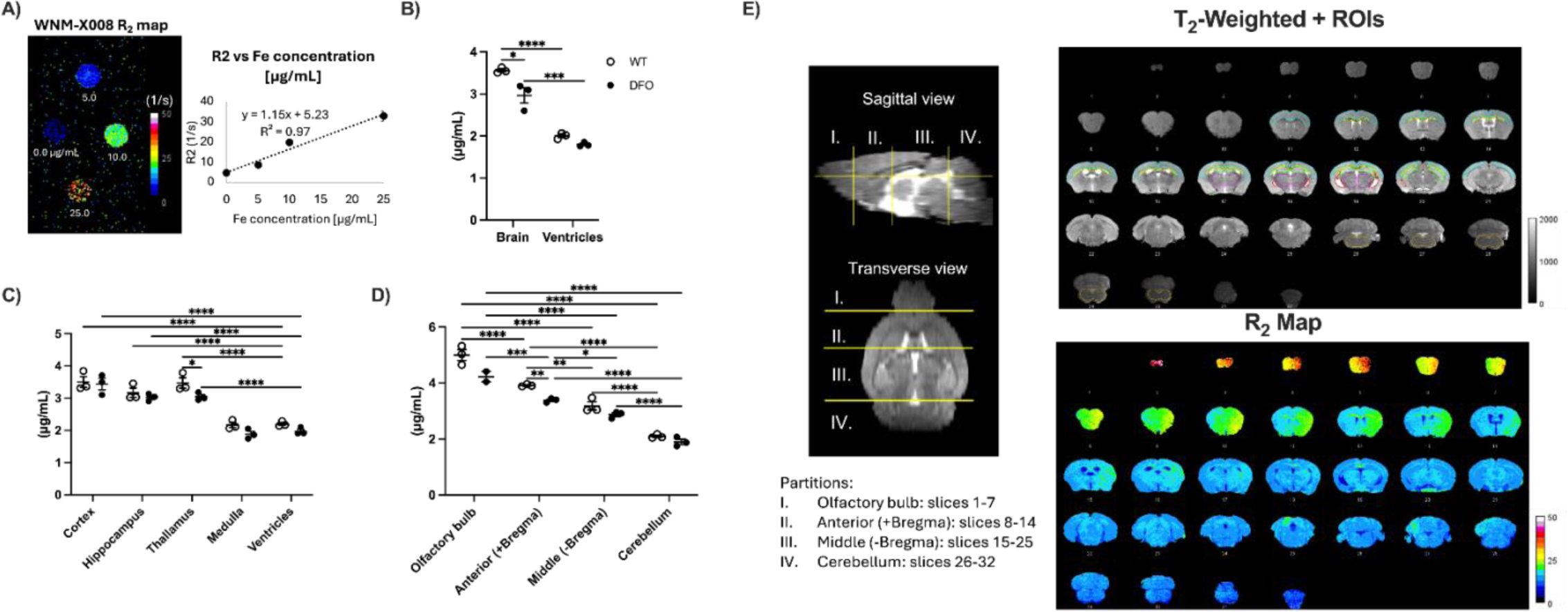
**A)** Standard curve for MRI using phantom containing iron oxide particles (ug/mL) as a reference for iron levels by MRI. Iron concentration was interpolated from R_2_ maps for **B)** whole brain and ventricles, **C)** defined regions of interest (ROI) or **D)** by section. **E)** ROI were manually and independently delineated by 2 experimenters, including the cortical mantle (cyan), hippocampus (green), thalamus (magenta), white matter (yellow), lateral ventricles (red), and medulla (orange). Partitions were categorized by slice position: I. Olfactory bulb (slices 1-7), II. Anterior (+Bregma, slices 8-14), III. Middle (-Bregma, slices 15-25), and IV. Cerebellum (slices 26-32). R_2_ maps were generated from the MEMS sequences on the same coronal slices. R_2_ calibration bars are in units of 1/s. Mean R_2_ were obtained from the slices that included the ROIs. Mean R_2_ values (circles) were obtained from segmented brain matter and ventricle regions (n=3/group), averaging R_2_ values across all brain slices in brain matter and ventricle regions. Mice were scanned at the same position in the MRI scanner. Two-way ANOVA with Tukey’s posthoc: *p<0.05, **p<0.01, ***p<0.001, ****p<0.0001.

**Supplemental Figure 2:**
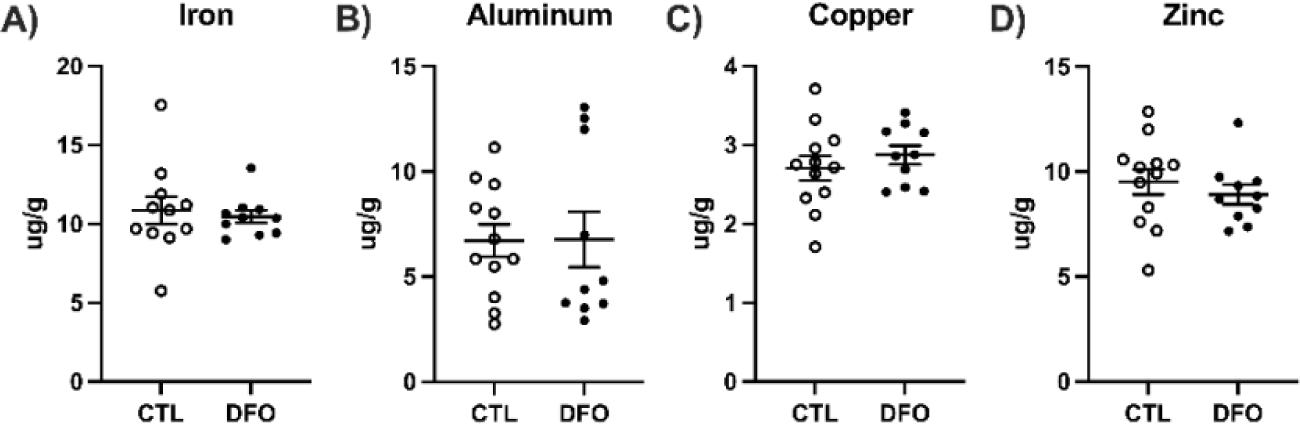
DFO treatment did not alter other metals in cortex. ICP-MS for total **A)** iron, **B)** aluminum, **C)** copper, and **D)** zinc.

**Supplemental Figure 3:**
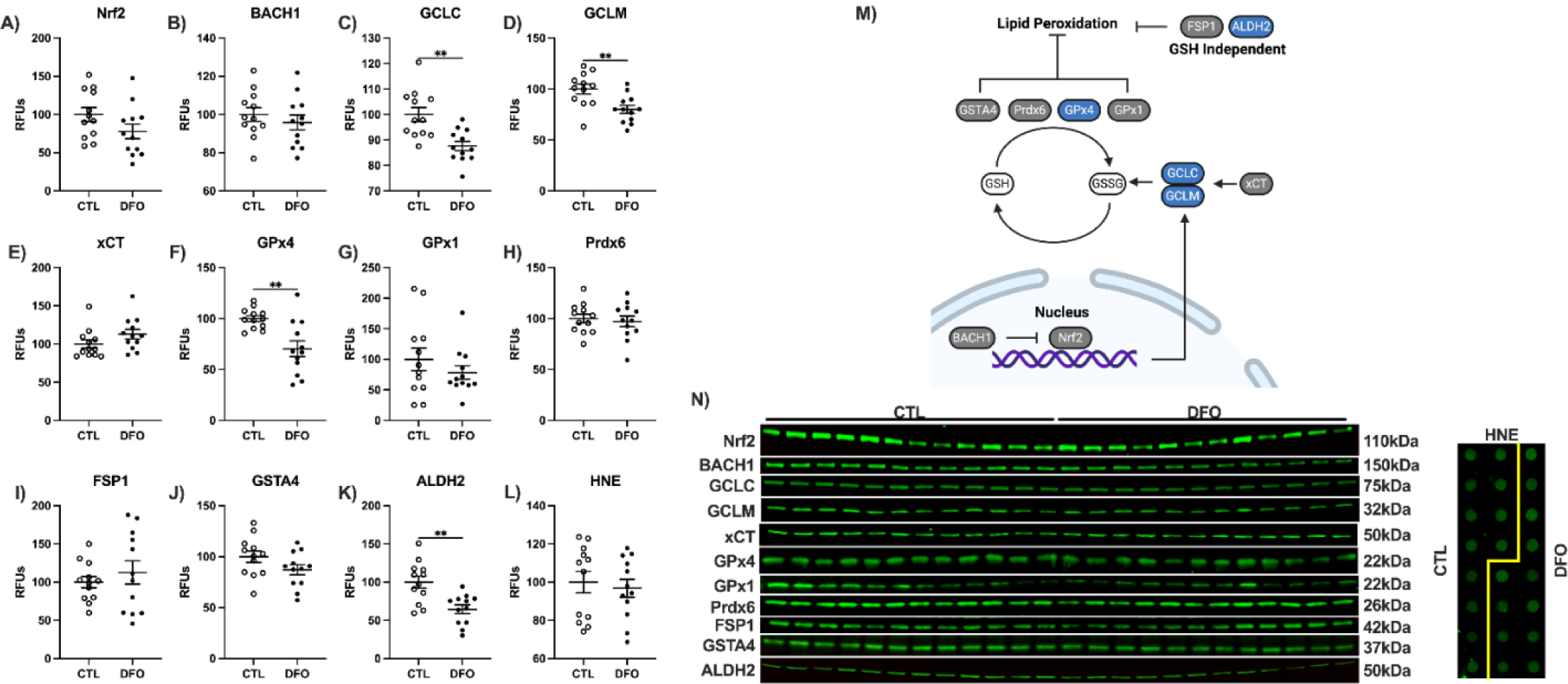
Antioxidant signaling is minimally altered by DFO. Western blots, relative fluorescent units (RFUs) for **A)** Nrf2, **B)** BACH1, **C)** GCLC, **D)** GCLM, **E)** xCT, **F)** GPx4, **G)** GPx1, **H)** Prdx6, **I)** FSP1, **J)** GSTA4, **K)** ALDH2, and dot blot for **L)** HNE. **M)** Schematic representation of antioxidant signaling with DFO treatment relevant to lipid peroxidation. Blue, increase; grey, no change; two-tailed t-test: **p<0.01.

## DISCUSSION

The iron chelator DFO cognitively benefited AD in its first and only trial in 1991 (15). To explore potential mechanisms, the present study evaluated brain responses in wildtype (WT) C7BL/6 mice. DFO decreased brain iron by 18% (MRI) and decreased seven major proteins that mediate iron metabolism including the iron storage proteins ferritin light and heavy chain by at least 25%. APP and secretase enzymes also decreased by 30%. Thus, DFO alters multiple biochemical processes related to AD in young WT mice that include amyloid processing enzymes, APP itself, and antioxidant repair. Potential DFO treatment for early-stage AD by DFO should consider the benefits of lowered APP and secretase enzymes.

We also recently analyzed DFO responses of the ApoE Familial AD (EFAD) mouse model, which decreased iron-mediated lipid peroxidation through antioxidant signaling, as well as decreasing fibrillar amyloid (4). Because these EFAD mice have weaker blood-brain barrier BBB integrity (38), and ApoE alleles influence BBB integrity (39), it was important to know how WT mice respond to DFO. In both studies, dietary DFO altered brain iron signaling with less impact on WT for antioxidant pathways than EFAD mice.

WT mice showed brain region specificity for decreased iron, using a novel MRI assay to show decreased iron only in thalamus and olfactory bulb. The thalamus has more BBB permeability and is dorsal to the hippocampus, making it a potential site for drug delivery AD treatments (40). In comparison AD vulnerable regions such as the frontal cortex and hippocampus have less BBB permeability associated with pericyte coverage (26).

Despite lack of DFO impact on cerebral cortex iron levels, DFO decreased levels of iron signaling proteins that mediate import, export, and storage. Since DFO treatment lowered iron levels under physiological levels in whole brain, the cerebral cortex responses to DFO for decreased iron signaling proteins suggest iron retention. These WT findings differed in direction from responses of EFAD mice, in which DFO increased iron signaling proteins; iron was not assayed in EFAD mice. Additionally, antioxidant signaling increased in EFAD and decreased in WT with DFO. The mouse genotype difference in response of iron signaling and antioxidant pathways may also be a result of differences in BBB integrity or the potentially higher baseline of iron due to the documented MBs in EFAD mice (5). These MBs release red blood cells which lyse readily by oxidation, releasing hemoglobin (41). The breakdown of hemoglobin yields extracellular hemosiderin deposits, which forms at sites of MBs (42–44) that contain 70-fold more extracellular than intracellular iron (4). In AD, these iron depots may increase lipid peroxidation by Fenton chemistry (2), which further damages proteins, DNA, and lipids. HNE, one major product of lipid peroxidation did not change in WT mice, which is consistent with their lower oxidative burden. In EFAD mice, DFO treatment for the same duration decreased HNE levels, which further substantiates its potential use for the treatment of AD. Antioxidants responsible for mitigating iron mediated oxidative damage such as HNE were largely unaltered with DFO treatment. This is further supported by the reductions by DFO for GSH biosynthesis proteins and GPx4 which reduces oxidized phospholipids and sterols. EFAD mice had increased Nrf2, GPx4, and GCLC, further supporting the potential of iron chelation for AD.

Iron and other transition state metals mediate the regulation of APP protein levels. Iron regulatory proteins dictate the fate of APP protein by binding to its mRNA facilitating or inhibiting translation (45). Low concentrations of iron are associated with inhibition of APP translation consistent with the decrease of APP protein in DFO treated cortex. The secretase enzymes BACE1 and PSEN1 that cleave APP for its Aβ peptides followed suit. BACE1 protein is dependent on copper levels, also chelated by DFO (46). Binding of BACE1 and APP is promoted by iron (47), further connecting regulation of these proteins to concentration of iron and other transition state metals. While ADAM10 was not significantly decreased by DFO, the downward trend was consistent with iron-overload diseases (48). ApoE and LRP1, which have roles in Aβ clearance, are regulated by iron concentrations at the mRNA and protein level in both neurons and astrocytes (49). These findings further support DFO’s capacity to modify iron sensitive signaling in brain tissues.

In summary, we show that acute DFO treatment on WT mice decreased cerebral cortex APP levels, its related secretase enzymes, and reduced brain iron storage protein. Limitations include the potential confounds of additional transition state metals scavenged by DFO and other iron chelators which are clearly involved in the regulation of APP processing, and the bulk analysis of protein changes to DFO by all brain cell types. Other confounds include BBB differences where rodents have similar BBB penetrance (50) which are less permeable than humans (51). These differences along with ApoE alleles which influence BBB integrity (39) may influence dosages given. Future studies may determine optimal duration and dosage levels for AD which will vary by neuropathological burden. Regardless, these data give a strong first basis for the potential of non-invasive treatment of DFO for AD or other neuropathologies.

## METHODS

### Mice

C57BL/6 mice from Jackson laboratories were housed by the University of Southern California Department of Animal Resources. Mice (n=12/group; equal sex) had ad libitum access to water and standard or DFO lab chow made by Research diets inc. (New Brunswick, NJ). Mice were housed in groups of 5 at 22°C/ 30% humidity, with standard nesting material and light cycles of 0600-1800 hr. Studies were approved by the USC Animal Use and Care Committee (IACUC# 20417). DFO treatment was administered by diet in rodent chow (10mg/kg/day) for two weeks starting at 5.5-months of age. At 6-months mice were anesthetized with isoflurane, euthanized by cardiac puncture, and PBS perfused. Cortex was stored at -80°C until homogenization in RIPA buffer (Millipore, Burlington, MA) with protease and phosphatase inhibitor cocktails (ThermoFisher, Waltham, MA).

### MRI acquisition

All live animal scans were approved and performed at the Functional Biological Imaging Core at the Zilkha Neurogenetic Institute (University of Southern California). The cryogen-free MR Solutions PET/MRI 7T system was equipped with a bore size of ∼24 cm, a maximum gradient up to 600 mT/m, and a 20 mm internal diameter quadrature bird cage mouse head coil. Animals were prepared for scans by undergoing anesthesia induction at 1–1.5% isoflurane in room air in an induction chamber (SomnoSuite, Torrington, CT). The animal was then positioned onto a scanner bed (Minerve, Esternay, France) with a head contention, was temperature controlled, and was maintained at 36-37°C. Ophthalmic ointment was applied over the animal’s eyes. A pneumatic pillow was placed over the animal’s abdomen to monitor respiration during the scans (SMAII). During scans, the animal received 1.5-2% isoflurane in 95% oxygen from an oxygen concentrator, delivered via a nose cone. After the scans, animals were placed in a heated recovery chamber until they are ambulatory before returning them to their home cages.

A positioning gradient echo sequence was first acquired to prepare the slice stacks for the 2D multi-echo multi slice (MEMS) spin echo sequence. The MEMS parameters are as follows: echo times (TEs) = 12, 24, 36, 48, 60, 72, 84, 96, 108, and 120 ms; TR = 3366 ms; slice thickness = 0.5 mm; FOV = 20 mm x 20 mm; MS = 256 256; and NA = 1.

### MRI Analysis

T_2_ maps were generated from the MEMS images, through a pixel-by-pixel exponential fitting of the signal intensities across the different TE times. MATLAB Rocketship v.1.4 module was used to perform the parametric fits(53). All fits with an r^2^>0.6 were included. Using Fiji software, pixels with fits having r^2^ values <0.6 were set to not-a-number (NaN) and were not included in the analysis. Additionally, brain regions were extracted by manually delineating brain outlines on each slice and outside brain regions were set to NaN. Regions of interests (ROI) were manually delineated using the polygon tool in Fiji. R_2_ maps were generated from the T_2_ maps, using the relationship T_2_ = 1/R_2_. Mean R_2_ values for each ROI, for each subject, were obtained. GraphPad was used for planned comparisons between groups.

### Western and Blot

20 ug of protein was boiled at 75°C under denatured conditions and resolved on 4-20% gradient gels. Proteins were electroblotted using a Criterion blotter (Bio-Rad Laboratories, Hercules, CA) and transferred onto 0.45um polyvinyl difluoride membranes. Membranes were stained using Revert 700 fluorescent protein stain and imaged prior to blocking with LI-COR Intercept blocking buffer (LI-COR Biosciences, Lincoln, NE), followed by primary antibodies. Membranes incubated with IRDye 800CW and/or 700CW secondary antibodies and visualized by Odyssey (LI-COR Biosciences). Western blot data was quantified with ImageJ and normalized by total protein per lane.

### Dot Blot

20ug from RIPA lysate was loaded onto PVDF membranes by gravity filtration using a Bio-dot microfiltration apparatus (Bio-Rad Laboratories, Hercules, CA) for amyloid peptides and oxidative damage markers. Dot blots were handled as described for Western blots above.

### Biochemical Assays

RIPA lysates were analyzed for heme values quantified by Quantichrom assay (BioAssay Systems, Hayward, CA) per the manufacturer’s instructions.

### ICP-MS

20mg of cortex was cut with a ceramic scalpel and placed in metal free test tubes. Tissue was washed twice with PBS and homogenized with sterilized disposable plastic pestles in Chelex 100 (sigma, St. Louis, MO) with purified water (18.2 MΩ; Millipore, Burlington, MA). Homogenates were desiccated by vacuum centrifuge at 95°C for 90 min. and solubilized in trace metal free 70% HNO_3_ overnight. 30% H_2_O_2_ was then added followed by drying under heat. After resuspension in 2% HNO_3_, samples were analyzed by Agilent 7500ce ICP-MS in hydrogen mode with a practical detection limit of 10ppb and a relative standard deviation (RSD) of replicate measures between 0.2 to 5%. Total iron concentration was normalized to wet weight tissue.

### Statistics

GraphPad Prism 10 software (GraphPad, San Diego, CA) was used for graphing and statistical analysis. Comparisons of two groups were performed by two-tailed t-test or multiple groups by two-way ANOVA with Tukey’s HSD (p<0.05) with a confidence interval set at 95%. Data that were not normally distributed were analyzed by Mann Whitney U or Kruskal-Wallis with Dunn’s test.

## Acknowledgements & Funding

We thank Liying Zhao for performing ICP-MS measurements. Lab studies were supported by NIH grants to CEF (R01-AG051521, P50-AG05142, P01-AG055367) and Cure Alzheimer’s Fund.

## Ethics Declarations

All of the authors declare no competing interests.

## Data Availability

The authors declare that the data supporting the findings of this study are available within the paper and its Supplementary Information files. Should any raw data files be needed in another format they are available from the corresponding author upon reasonable request.

## Author Contributions

Max A. Thorwald (Conceptualization, Data curation; Formal Analysis; Investigation; Validation; Visualization; Writing – original draft; Writing – review & editing); Jose A. Godoy-Lugo (Conceptualization, Data curation; Formal Analysis; Investigation; Writing – review & editing); Naomi Sta Maria (Conceptualization, Data curation; Formal Analysis; Investigation; Methodology; Validation; Visualization; Resources; Software; Writing – original draft; Writing – review & editing); Ararat Chakhoyan (Data curation; Formal Analysis; Investigation; Methodology; Software; Validation; Writing – review & editing); Peggy O’Day (Data curation; Investigation; Resources; Methodology; Writing – review & editing); Russell E. Jacobs (Methodology; Writing – review & editing); Berislav Zlokovic (Conceptualization; Funding Acquisition; Resources; Supervision; Writing – review & editing); Caleb E. Finch (Conceptualization; Funding Acquisition; Project Administration; Resources; Supervision; Writing – original draft; Writing – review & editing).

## References

1. Butterfield DA, Lauderback CM. Lipid peroxidation and protein oxidation in Alzheimer’s disease brain: potential causes and consequences involving amyloid beta-peptide-associated free radical oxidative stress. Free Radic Biol Med. 2002 Jun 1;32(11):1050–60.

2. Montine TJ, Neely MD, Quinn JF, Beal MF, Markesbery WR, Roberts LJ, et al. Lipid peroxidation in aging brain and Alzheimer’s disease. Free Radic Biol Med. 2002 Sep 1;33(5):620–6.

3. Chen L, Dar NJ, Na R, McLane KD, Yoo K, Han X, et al. Enhanced defense against ferroptosis ameliorates cognitive impairment and reduces neurodegeneration in 5xFAD mice. Free Radical Biology and Medicine. 2022 Feb 20;180:1–12.

4. Thorwald MA, Godoy-Lugo JA, Garcia G, Silva J, Kim M, Christensen A, et al. Iron associated lipid peroxidation in Alzheimer’s disease is increased in lipid rafts with decreased ferroptosis suppressors, tested by chelation in mice [Internet]. bioRxiv; 2024 [cited 2024 Jun 22]. p. 2023.03.28.534324. Available

5. Cacciottolo M, Morgan TE, Finch CE. Age, sex, and cerebral microbleeds in EFAD Alzheimer disease mice. Neurobiol Aging. 2021 Jul;103:42–51.

6. van Bergen JMG, Li X, Hua J, Schreiner SJ, Steininger SC, Quevenco FC, et al. Colocalization of cerebral iron with Amyloid beta in Mild Cognitive Impairment. Sci Rep. 2016 Oct 17;6(1):35514.

7. Ayton S, Portbury S, Kalinowski P, Agarwal P, Diouf I, Schneider JA, et al. Regional brain iron associated with deterioration in Alzheimer’s disease: A large cohort study and theoretical significance. Alzheimer’s & Dementia. 2021;17(7):1244–56.

8. Damulina A, Pirpamer L, Soellradl M, Sackl M, Tinauer C, Hofer E, et al. Cross-sectional and Longitudinal Assessment of Brain Iron Level in Alzheimer Disease Using 3-T MRI. Radiology. 2020 Sep;296(3):619–26.

9. Ashraf A, Jeandriens J, Parkes HG, So PW. Iron dyshomeostasis, lipid peroxidation and perturbed expression of cystine/glutamate antiporter in Alzheimer’s disease: Evidence of ferroptosis. Redox Biol. 2020 May;32:101494.

10. Hernandez P, Johnson CA. Deferoxamine for aluminum toxicity in dialysis patients. ANNA J. 1990 Jun;17(3):224–8.

11. Farkas E, Csóka H, Micera G, Dessi A. Copper(II), nickel(II), zinc(II), and molybdenum(VI) complexes of desferrioxamine B in aqueous solution. Journal of Inorganic Biochemistry. 1997 Mar 1;65(4):281–6.

12. Farr AC, Xiong MP. Challenges and Opportunities of Deferoxamine Delivery for Treatment of Alzheimer’s Disease, Parkinson’s Disease, and Intracerebral Hemorrhage. Mol Pharm. 2021 Feb 1;18(2):593–609.

13. Olivieri NF, Brittenham GM. Iron-Chelating Therapy and the Treatment of Thalassemia. Blood. 1997 Feb 1;89(3):739–61.

14. Kang H, Han M, Xue J, Baek Y, Chang J, Hu S, et al. Renal clearable nanochelators for iron overload therapy. Nat Commun. 2019 Nov 13;10:5134.

15. Crapper McLachlan DR, Dalton AJ, Kruck TP, Bell MY, Smith WL, Kalow W, et al. Intramuscular desferrioxamine in patients with Alzheimer’s disease. Lancet. 1991 Jun 1;337(8753):1304–8.

16. Fujisawa K, Takami T, Matsumoto T, Yamamoto N, Yamasaki T, Sakaida I. An iron chelation-based combinatorial anticancer therapy comprising deferoxamine and a lactate excretion inhibitor inhibits the proliferation of cancer cells. Cancer Metab. 2022 May 12;10:8.

17. Keberle H. THE BIOCHEMISTRY OF DESFERRIOXAMINE AND ITS RELATION TO IRON METABOLISM. Ann N Y Acad Sci. 1964 Oct 7;119:758–68.

18. Ward RJ, Dexter D, Florence A, Aouad F, Hider R, Jenner P, et al. Brain iron in the ferrocene-loaded rat: its chelation and influence on dopamine metabolism. Biochem Pharmacol. 1995 Jun 16;49(12):1821–6.

19. Shachar DB, Kahana N, Kampel V, Warshawsky A, Youdim MBH. Neuroprotection by a novel brain permeable iron chelator, VK-28, against 6-hydroxydopamine lession in rats. Neuropharmacology. 2004 Feb;46(2):254–63.

20. Palmer C, Roberts RL. Deferoxamine Starch Conjugate Provides Partial Protection from Hypoxic-Ischemic Brain Injury in Neonatal Rats † 1749. Pediatr Res. 1997 Apr;41(4):294–294.

21. Hurn PD, Koehler RC, Blizzard KK, Traystman RJ. Deferoxamine reduces early metabolic failure associated with severe cerebral ischemic acidosis in dogs. Stroke. 1995 Apr;26(4):688–94; discussion 694-695.

22. Fine JM, Renner DB, Forsberg AC, Cameron RA, Galick BT, Le C, et al. Intranasal deferoxamine engages multiple pathways to decrease memory loss in the APP/PS1 model of amyloid accumulation. Neurosci Lett. 2015 Jan 1;584:362–7.

23. Zhang Y, He M lin. Deferoxamine enhances alternative activation of microglia and inhibits amyloid beta deposits in APP/PS1 mice. Brain Research. 2017 Dec 15;1677:86–92.

24. Guo Z, Lin J, Sun K, Guo J, Yao X, Wang G, et al. Deferoxamine Alleviates Osteoarthritis by Inhibiting Chondrocyte Ferroptosis and Activating the Nrf2 Pathway. Front Pharmacol. 2022 Mar 14;13:791376.

25. Inoue H, Hanawa N, Katsumata SI, Katsumata-Tsuboi R, Takahashi N, Uehara M. Iron deficiency induces autophagy and activates Nrf2 signal through modulating p62/SQSTM. Biomed Res. 2017;38(6):343–50.

26. Winkler EA, Sengillo JD, Bell RD, Wang J, Zlokovic BV. Blood–spinal cord barrier pericyte reductions contribute to increased capillary permeability. J Cereb Blood Flow Metab. 2012 Oct;32(10):1841–52.

27. Bauminger ER, Harrison PM, Hechel D, Nowik I, Treffry A. Iron (III) can be transferred between ferritin molecules. Proc Biol Sci. 1991 Jun 22;244(1311):211–7.

28. Mancias JD, Wang X, Gygi SP, Harper JW, Kimmelman AC. Quantitative proteomics identifies NCOA4 as the cargo receptor mediating ferritinophagy. Nature. 2014 May;509(7498):105–9.

29. Rogers JT, Randall JD, Cahill CM, Eder PS, Huang X, Gunshin H, et al. An iron-responsive element type II in the 5’-untranslated region of the Alzheimer’s amyloid precursor protein transcript. J Biol Chem. 2002 Nov 22;277(47):45518–28.

30. Igarashi K, Hoshino H, Muto A, Suwabe N, Nishikawa S, Nakauchi H, et al. Multivalent DNA Binding Complex Generated by Small Maf and Bach1 as a Possible Biochemical Basis for β-Globin Locus Control Region Complex *. Journal of Biological Chemistry. 1998 May 8;273(19):11783–90.

31. Venugopal R, Jaiswal AK. Nrf1 and Nrf2 positively and c-Fos and Fra1 negatively regulate the human antioxidant response element-mediated expression of NAD(P)H:quinone oxidoreductase1 gene. Proc Natl Acad Sci U S A. 1996 Dec 10;93(25):14960–5.

32. Chen Y, Shertzer HG, Schneider SN, Nebert DW, Dalton TP. Glutamate cysteine ligase catalysis: dependence on ATP and modifier subunit for regulation of tissue glutathione levels. J Biol Chem. 2005 Oct 7;280(40):33766–74.

33. Thomas JP, Geiger PG, Maiorino M, Ursini F, Girotti AW. Enzymatic reduction of phospholipid and cholesterol hydroperoxides in artificial bilayers and lipoproteins. Biochimica et Biophysica Acta (BBA) - Lipids and Lipid Metabolism. 1990 Aug 6;1045(3):252–60.

34. Fisher AB. Peroxiredoxin 6 in the repair of peroxidized cell membranes and cell signaling. Arch Biochem Biophys. 2017 Mar 1;617:68–83.

35. Castro JP, Jung T, Grune T, Siems W. 4-Hydroxynonenal (HNE) modified proteins in metabolic diseases. Free Radical Biology and Medicine. 2017 Oct 1;111:309–15.

36. Du K, Oh SH, Dutta RK, Sun T, Yang WH, Chi JTA, et al. Inhibiting xCT/SLC7A11 induces ferroptosis of myofibroblastic hepatic stellate cells but exacerbates chronic liver injury. Liver Int. 2021 Sep;41(9):2214– 27.

37. Tang D, Chen X, Kang R, Kroemer G. Ferroptosis: molecular mechanisms and health implications. Cell Res. 2021 Feb;31(2):107–25.

38. Montagne A, Nikolakopoulou AM, Huuskonen MT, Sagare AP, Lawson EJ, Lazic D, et al. APOE4 accelerates advanced-stage vascular and neurodegenerative disorder in old Alzheimer’s mice via cyclophilin A independently of amyloid-β. Nat Aging. 2021 Jun;1(6):506–20.

39. Bell RD, Winkler EA, Singh I, Sagare AP, Deane R, Wu Z, et al. Apolipoprotein E controls cerebrovascular integrity via cyclophilin A. Nature. 2012 May 16;485(7399):512–6.

40. Huh H, Park TY, Seo H, Han M, Jung B, Choi HJ, et al. A local difference in blood-brain barrier permeability in the caudate putamen and thalamus of a rat brain induced by focused ultrasound. Sci Rep. 2020 Nov 6;10(1):19286.

41. Chang R, Castillo J, Zambon AC, Krasieva TB, Fisher MJ, Sumbria RK. Brain Endothelial Erythrophagocytosis and Hemoglobin Transmigration Across Brain Endothelium: Implications for Pathogenesis of Cerebral Microbleeds. Front Cell Neurosci. 2018 Sep 6;12:279.

42. O’Connell MJ, Ward RJ, Baum H, Peters TJ. The role of iron in ferritin- and haemosiderin-mediated lipid peroxidation in liposomes. Biochem J. 1985 Jul 1;229(1):135–9.

43. Litwack G. Chapter 19 - Micronutrients (Metals and Iodine). In: Litwack G, editor. Human Biochemistry (Second Edition) [Internet]. Boston: Academic Press; 2022 [cited 2022 May 19]. p. 647–701. Available from: https://www.sciencedirect.com/science/article/pii/B9780323857185000182

44. Janaway BM, Simpson JE, Hoggard N, Highley JR, Forster G, Drew D, et al. Brain haemosiderin in older people: pathological evidence for an ischaemic origin of magnetic resonance imaging (MRI) microbleeds. Neuropathol Appl Neurobiol. 2014 Apr;40(3):258–69.

45. Khan MA, Mohammad T, Malik A, Hassan MI, Domashevskiy AV. Iron response elements (IREs)-mRNA of Alzheimer’s amyloid precursor protein binding to iron regulatory protein (IRP1): a combined molecular docking and spectroscopic approach. Sci Rep. 2023 Mar 28;13(1):1–17.

46. Lin R, Chen X, Li W, Han Y, Liu P, Pi R. Exposure to metal ions regulates mRNA levels of APP and BACE1 in PC12 cells: Blockage by curcumin. Neuroscience Letters. 2008 Aug 8;440(3):344–7.

47. Gong L, Tian X, Zhou J, Dong Q, Tan Y, Lu Y, et al. Iron Dyshomeostasis Induces Binding of APP to BACE1 for Amyloid Pathology, and Impairs APP/Fpn1 Complex in Microglia: Implication in Pathogenesis of Cerebral Microbleeds. Cell Transplant. 2019 Aug;28(8):1009–17.

48. Maras JS, Das S, Sharma S, Sukriti S, Kumar J, Vyas AK, et al. Iron-Overload triggers ADAM-17 mediated inflammation in Severe Alcoholic Hepatitis. Sci Rep. 2018 Jul 6;8(1):10264.

49. Xu H, Perreau VM, Dent KA, Bush AI, Finkelstein DI, Adlard PA. Iron Regulates Apolipoprotein E Expression and Secretion in Neurons and Astrocytes. J Alzheimers Dis. 2016;51(2):471–87.

50. Murakami H, Takanaga H, Matsuo H, Ohtani H, Sawada Y. Comparison of blood-brain barrier permeability in mice and rats using in situ brain perfusion technique. Am J Physiol Heart Circ Physiol. 2000 Sep;279(3):H1022–1028.

51. Syvänen S, Lindhe O, Palner M, Kornum BR, Rahman O, Långström B, et al. Species differences in blood-brain barrier transport of three positron emission tomography radioligands with emphasis on P-glycoprotein transport. Drug Metab Dispos. 2009 Mar;37(3):635–43.

52. Zhang Y, Brady M, Smith S. Segmentation of brain MR images through a hidden Markov random field model and the expectation-maximization algorithm. IEEE Trans Med Imaging. 2001 Jan;20(1):45–57.

53. Barnes SR, Ng TSC, Santa-Maria N, Montagne A, Zlokovic BV, Jacobs RE. ROCKETSHIP: a flexible and modular software tool for the planning, processing and analysis of dynamic MRI studies. BMC Medical Imaging. 2015 Jun 16;15(1):19.

